# Winter flowering by keystone desert perennials explained using climate change attribution

**DOI:** 10.64898/2026.05.22.727294

**Authors:** Jeremy B. Yoder, Colin J. Carlson, Christopher W. Callahan

## Abstract

Climate change is expected to touch every ecological community on the planet, but connecting climate change to specific ecological events remains challenging. The emerging practice of climate change attribution can identify how anthropogenic climate change contributes to extreme weather events—but has not yet been applied to unusual ecological events. Here, we demonstrate attribution of a recent, striking ecological anomaly: Joshua trees (*Yucca brevifolia* and *Y. jaegeriana*) flowering in October 2025, fully four months earlier than normal. We used crowdsourced records of Joshua tree flowering to train a machine learning model that recovers weather triggers of flowering, and successfully predicts regular-seasonal and out-of-season flowering events. We then simulated weather without human-caused climate change using the output of 10 global climate models, and projected Joshua tree flowering under those counterfactual conditions with our trained model. Surprisingly, we found out-of-season blooms were driven by high winter rainfall, not rising temperatures—and therefore, are probably the result of natural weather variability. Our results place out-of-season flowering in context with climate change threats facing Joshua trees, while providing a prototype attribution analysis of an extreme ecological event and identifying priorities for future climate modeling.

## Introduction

Anthropogenic climate change has the potential to alter ecological processes in every biome, and many such alterations are already apparent (Parmesan et al., 2022; Parmesan and Yohe, 2003). The ecological effects of climate change generally scale continuously with warming average temperatures, but climate change may also manifest as extreme events such as droughts, heat waves, and storms (Seneviratne and Zhang, 2021). More frequent extreme weather events should logically have ecological impacts, creating anomalies like irruptions, population booms, and mass die-offs (Bai et al., 2024; Baum et al., 2026; Stager et al., 2026). However, linking specific anomalous ecological events to global climate trends is not straightforward.

Recent years have seen substantial advances in the attribution of specific extreme events to human-caused climate change (NASEM, 2016). The standard approach is to estimate the probability or intensity of those events under factual (i.e., “historical” climates: history as it happened, including human influence on the Earth system) and counterfactual (“natural” climates: simulations that ideally include natural weather variability, but exclude human influence) scenarios (Swain et al., 2020). These methods have established beyond a reasonable doubt that climate change has increased the frequency and intensity of certain events, such as heat waves (IPCC, 2021). However, for phenomena such as shifts in seasonal rainfall averages or variability, the signal of climate change is much less clear, and these changes are often dominated by natural variability (Lehner et al., 2018). There is growing interest in applying attribution science to the human and ecological impacts of climate change (Carlson et al., 2024, 2025; Dudney et al., 2025), but the complexity of biological responses to local climate and the limited availability of long-term ecological data often prevent confident attribution to climate change (Parmesan et al., 2011; Zwiers and Hegerl, 2008).

Here, we ask whether a specific anomalous ecological event can be attributed to climate change: in 2025, Joshua trees (Fig. 1A; *Yucca brevifolia* Engelm. and *Y. jaegeriana* McKelvey, Asparagaceae) began flowering in October, roughly four months in advance of their typical flowering onset (Fig. 1B).

**Fig. 1.**
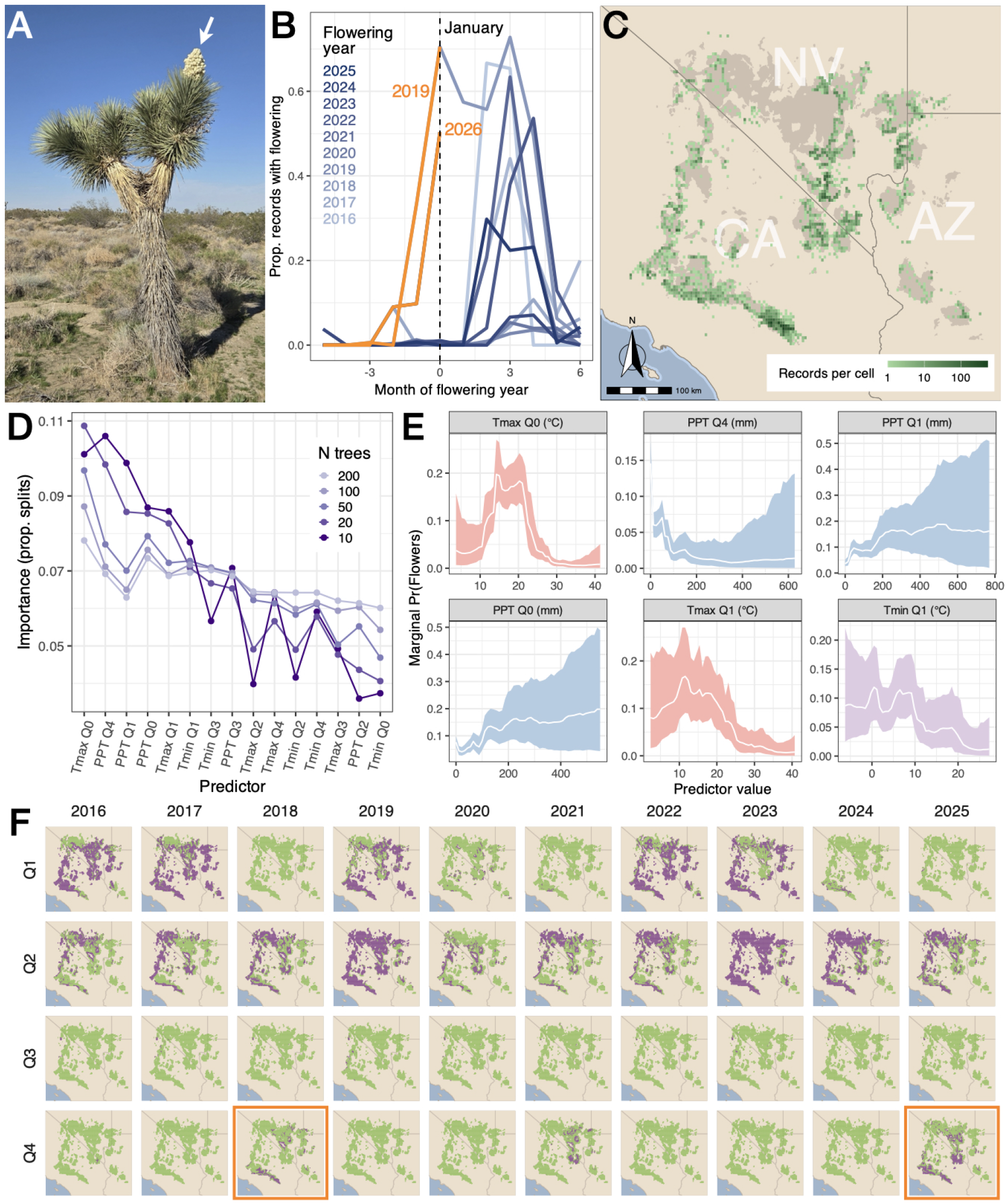
Crowdsourced data tracks anomalous out-of-season flowering of Joshua trees and provides a basis to train BART models that recover both regular and anomalous flowering. (A) A western Joshua tree (*Yucca brevifolia*) with open flowers on 2 Nov 2025, from iNaturalist record 324625796, observed by contributor Drew Kaiser (username `drew_kaiser`). (B) Proportion of iNaturalist records indicating the presence of flower buds or open flowers, by month, for 2016–2025; lines for more recent years are darker, and line segments for observed flowering in the months leading up to the 2019 and 2026 flowering years (i.e., late 2018 and 2025) are highlighted in orange. (C) Geographic distribution of iNaturalist records of Joshua trees, aggregated to a ∼4km raster grid (darker cells, more records) overlaid on Joshua trees’ range (khaki shading; Esque et al. 2023) across southern California, Nevada, Utah, and Arizona, USA. (D) Predictor importance plot for RI-BART models of varied complexity (number of trees) predicting quarterly Joshua tree flowering. (E) Posterior partial effects of the six most informative weather predictors of Joshua tree flowering (median, white lines; colored areas, 95% density interval). (F) Regions in the Joshua tree range with flowering (purple) or no flowering (green) predicted by the final working RI-BART model, for each quarter of 2016–2025; anomalous flowering events in the fourth quarters of 2018 and 2025 highlighted by orange boxes.

Joshua trees are long-lived arborescent monocots, endemic to the Mojave Desert of the southwestern United States. They are foundational members of multiple Mojave plant communities, providing food and shelter for a wide variety of native wildlife (Peattie, 1950; Terrill et al., 2019; Yoder and Smith, 2024), and they have attracted increasing attention as “umbrella species” for this endemic biodiversity, as they face mounting threats from habitat loss to development, changing wildfire regimes, and climate change (Smith et al., 2023). The trees only set seed when pollinated by highly specialized yucca moths (genus *Tegeticula*; Godsoe et al. 2008; Pellmyr 2003), and the prospect that this mutualism could be disrupted by climate change-driven phenological shifts is a longstanding conservation concern (Smith et al., 2023).

Joshua trees follow a masting reproductive strategy that tracks inter-annual variation in precipitation. We recently modeled variation in Joshua tree flowering driven by annual variation in temperature and precipitation, and found that the frequency of masting years has likely increased slightly since the early 20th century, as annual precipitation has become more variable (Yoder et al., 2024). This finding, and prior speculation about the causes of a similar out-of-season flowering event in autumn of 2018 (Brenskelle et al., 2021), led naturally to the hypothesis that the 2025 flowering anomaly was the result of human-caused climate change (Kuta, 2026; Rode, 2026).

We used the excellent resource for tracking Joshua tree flowering activity provided by the iNaturalist crowdsourcing platform (inaturalist.org) to model regular-seasonal and out-of-season flowering from 2016 to 2025, and to infer whether anomalous flowering is due to recent climate change. Our model identified precipitation as a key driver of both regular and out-of-season flowering, and recovered regular and out-of-season flowering events within the time-frame of our training data, as well as flowering recorded in independent records as early as 1913. We used this model to simulate flowering under a counter-factual scenario in which the climate change signal has been removed from local temperature and precipitation, based on projections of 10 global climate model simulations—and we found no significant difference in predicted regular or out-of-season flowering activity. We conclude that observed out-of-season flowering events are not likely to be due to global climate trends, but may instead be a response to low-frequency early winter precipitation events.

## Methods

We conducted all analyses in R (version 4.5, R Core Team 2025), with specialized functions as indicated.

### Annotation and compilation of crowdsourced records

We obtained records of *Yucca brevifolia* and *Y. jaegeriana* flowering from iNaturalist (inaturalist.org), a community science platform where contributors upload photos of organisms with the time, date, and location of their observations (**?**). In our prior study modeling Joshua tree masting activity (Yoder et al., 2024), we confirmed species identity and annotated presence of flower buds, open flowers, fruits, or no evidence of flowering from images submitted with thousands of iNaturalist observations, using features of the iNaturalist online interface. As the 2025 anomalous flowering event emerged, we solicited observations of flowering Joshua trees through a “project” page within the iNaturalist interface to organize contributors’ efforts. We used functions from the package rinat to access the iNaturalist database through its API. We queried for “Research Grade” records, which have species identification confirmed by at least two iNaturalist contributors with no contradictory identifications, from the years 2008 (when iNaturalist launched) through 2025, and further filtered downloaded records to remove any with location uncertainty greater than 2km. Years prior to 2016 had dramatically poorer sampling as a consequence of the smaller iNaturalist contributor community at that time (Yoder et al., 2024), so we conducted downstream analysis using records from 2016 through 2025.

### Linking flowering records to weather data

We obtained monthly weather data from PRISM (PRISM Climate Group, Oregon State University, 2014), which provides spatially interpolated raster-formatted records of temperature (minimum and maximum, in degrees Centigrade), vapor pressure deficit (minimum and maximum, in hPa) and total precipitation (in millimeters) across the contiguous U.S. from 1895 to the present. For analysis, we summarized monthly records into calendar quarters: January–March, April–June, July–September, and October–December.

To link flowering observed in the iNaturalist records to quarterly weather data, we aggregated the iNaturalist records by year, quarter, and grid cell within the ∼4km raster grid of the PRISM data layers. Of the records within each grid cell in a given year and quarter, we determined whether any indicated flowering (annotated with flowers budding, or open flowers), and if so, we marked that grid cell as having *flowering present* in that year and quarter; otherwise we marked it as having *flowering absent*. We matched each observation of quarterly flowering presence/absence with the corresponding values of maximum and minimum temperature (°C) and total precipitation (mm) for each of five quarters, counting backwards from the quarter in which flowering was observed: Q0 for the quarter of observation, Q1 for the quarter immediately prior, and so on back to Q4, the same quarter as Q0 but in the prior year. (So, e.g., for an observation in April of 2019, Q0 would be April– June 2019, Q1 would be January–March of 2019, Q2 would be October–December 2018, Q3 would be July–September 2018, and Q4 would be April– June 2018.)

### Modeling flowering activity

We trained Bayesian additive regression tree (BART) models predicting flowering activity in a given year, quarter, and location with temperature and precipitation in the quarter in which flowering was observed (Q0) and the four quarters prior (Q1–Q4) as defined above. BART models are sum-of-tree classification and regression tree machine-learning models, which have notable advantages for applications in ecology (Carlson, 2020; Carlson et al., 2022; Chipman et al., 2010). In particular, a BART variant, RIBART, allows inclusion of a random-intercept effect added to the prediction of the main model to test for confounding differences due to stratification of the training data (Carlson et al., 2022).

We used functions in the `dbarts` (Dorie, 2023) and `embarcadero` (Carlson, 2020) packages to train and evaluate BART models of flowering—i.e., models predicting the presence or absence of flowering in a given year and quarter. We specifically used RI-BART models for primary analysis, with quarter of the year as the RI effect, to control for anticipated confounding between seasonal variation in our weather predictors and the regular flowering season. We identified informative predictors using a predictor selection procedure provided by `embarcadero`, which identifies the strongest predictors by training RI-BART models with 10, 20, 50, 100, or 200 component trees. Predictors that have consistently higher frequency of inclusion in simpler models (i.e., models with fewer trees) are the most informative predictors (Chipman et al., 2010). After predictor selection, we trained a standard RIBART model with only the top predictors, plus the RI effect of quarter, for downstream analyses.

To evaluate the model’s performance, we used it to predict quarterly flowering activity from the PRISM archive records for each year 1900–2025. We directly examined the spatial extent of predicted quarterly flowering for the years in our 10-year training data time-frame (2016–2025, Fig. 1F), and we validated predictions in other years against independent observations of Joshua tree flowering from the National Phenology Network database (USA-NPN, USA National Phenology Network 2026) and digitized herbarium records (California Consortium of Herbaria, 2024; Texas and Oklahoma Regional Consortium of Herbaria, 2024).

### Flowering predicted under counterfactual conditions

To test the hypothesis that the anomalous out-of-season flowering events in the final quarters of 2018 and 2025 were attributable to global climate change, we simulated temperature and precipitation for each year 2016–2025 under counterfactual conditions without recent climate change. To isolate the signal of climate change, we compared monthly temperature and precipitation from the “historical” and “historical-nat” (natural) simulations from 10 global climate models (GCMs) from the Sixth Phase of the Coupled Model Intercomparison Project (CMIP6) (Eyring et al., 2016; Gillett et al., 2016). The historical simulations are run with all observed forcings including greenhouse gas emissions (GHGs), and the natural simulations are run with natural forcings such as solar and volcanic changes but without effects of GHGs or land use change, allowing us to isolate the anthropogenic climate change signal. We interpolated each simulation to the PRISM grid and took the difference between the historical and natural simulations separately for each GCM to yield the signal of climate change, or “delta” (Figs. S3,4).

We subtracted these climate change “deltas” from the observed weather records to yield “counterfactual” weather that includes realistic year-to-year variability but no signal of anthropogenic climate change, following standard practices in the literature (e.g., Diffenbaugh et al., 2021). We then used these counterfactual weather records to predict flowering with the working model. We compared flowering predicted from the counterfactual weather records to predictions from actual records in terms of the proportion of the Joshua tree range (khaki shading, Fig. 1C) with flowering predicted in each quarter for the years 2016–2025.

## Results

### iNaturalist records capture out-of-season flowering events

The iNaturalist database provided 13,874 records of Joshua tree flowering for 2016– 2025. Organizing these records by month of flowering year, the out-of-season blooms in late 2018 and 2025 are qualitatively different from variation in flowering onset seen in other years, as months-early initiation of the 2019 and 2026 flowering years (Fig. 1B, orange lines). Aggregating the records to the raster grid of PRISM weather data and by quarter, we obtained 5,070 quarterly observations of flowering presence or absence, covering the breadth of the geographic distribution for both Joshua tree species (Fig. 1C).

### Machine learning recovers climatic triggers of both in- and out-of-season flowering activity

Predictor selection for random intercept Bayesian additive regression tree (RI-BART) models with quarter as the RI effect identified six top predictors of quarterly Joshua tree flowering (Fig. 1D): Maximum temperature and total precipitation in Q0 (quarter of flowering), maximum and minimum temperature in Q1 (quarter prior to flowering), and total precipitation in Q1 and Q4 (four quarters prior to flowering). Our working RI-BART model trained with these six predictors and a RI effect of calendar quarter had AUC = 0.86 for the training data (Fig. S1).

The posterior partial effects for the six weather predictors (Fig. 1E) indicate higher probability of flowering in quarters with Q0 and Q1 maximum temperature between 10 and 22 °C, greater Q0 and Q1 total precipitation but lower Q4 precipitation, and Q1 minimum temperature below 10 °C. In sum, these partial effects mean that the model finds greater probability of flowering in a wetter, cooler quarter that immediately follows a wetter, cooler quarter; and which is wetter than the same quarter in the previous year. The RI effect of calendar quarter acts to reduce the probability of flowering in the third and fourth quarter of a calendar year, reflecting lower probability of flowering outside the regular flowering season (Fig. S2).

Predicting quarterly flowering across the Joshua tree range while controlling for the quarterly RI effect, the model recovered out-of-season flowering events in the fourth quarters of 2018 and 2025 (Fig. 1F, orange boxes). This demonstrates that these events are predictable based on the same weather triggers as regular-season flowering.

For further validation, we obtained 453 independent records of quarterly Joshua tree flowering activity, from a much wider time-frame than our training data. The USA-National Phenology Network (USA National Phenology Network, 2026) provided 362 records spanning 2011–2025, and digitized herbarium records (California Consortium of Herbaria, 2024; Texas and Oklahoma Regional Consortium of Herbaria, 2024) provided 91 specimens spanning 1913–2021. Our model predicted significantly higher probability of flowering in quarters and locations for which these validation records indicated Joshua tree flowering (Welch’s two-sample t-test with *d f* = 341.5, *p* < 10^*−*6^), and had AUC = 0.81 for predictions of flowering in quarters and locations covered by the validation records.

### No estimated contribution of human-caused climate change to out-of-season flowering events

We used the model to project the proportion of the Joshua tree range with flowering in each quarter of 2016–2025, based on both actual (Fig. 2A,B) and simulated counterfactual weather for those years. Projected flowering based on actual and counterfactual weather had substantially overlapping 95% posterior intervals in all quarters (Fig. 2D), and the difference between projections from counterfactual weather and projections from actual weather was correspondingly small. Only the third quarter of 2019 and the third quarter of 2020 had factual-counterfactual differences significantly different from zero, and those differences amounted to a few percentage points. Neither of the quarters with anomalous flowering had factual-counterfactual differences significantly different from zero (Fig. 2C,E, blue-shaded quarters; Fig. S5). Thus, we find that human-caused climate change did not make a measurable contribution to the 2018 or 2025 flowering anomalies, which were instead driven by natural precipitation variability.

**Fig. 2.**
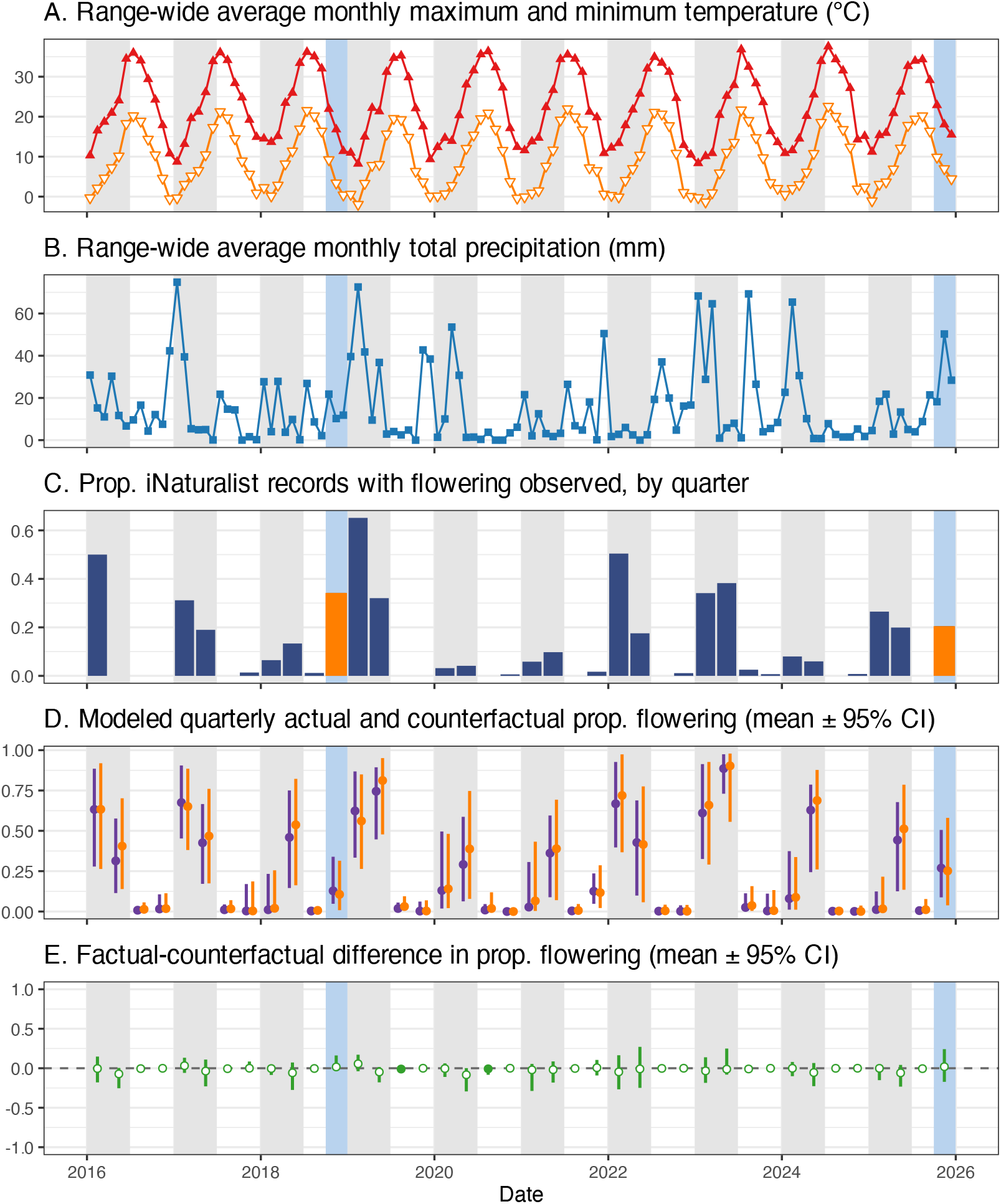
Weather, observed Joshua tree flowering, and flowering predicted by our model under factual and counterfactual conditions over the 2016–2025 study period. (A) Monthly maximum (red, upward-pointing triangles) and minimum temperature (orange, downward-pointing triangles), averaged over the Joshua tree species range in Fig. 1C (Esque et al., 2023). (B) Monthly total precipitation, averaged over the Joshua tree range. (C) Proportion of iNaturalist records with evidence of flower buds or open flowers, by quarter, with orange bars indicating anomalous flowering events. (D) Proportion of the Joshua tree range with flowering predicted in each quarter, using our final RI-BART model projecting from actual weather conditions (purple) or counterfactual conditions (orange); points give the mean and lines give the 95% posterior interval. (E) Counterfactual deviation (i.e., difference between predictions from actual and counterfactual weather) for the proportion of the range with predicted flowering in each quarter; points give the mean and lines give the 95% posterior interval as in (D), and points are filled if the posterior interval does not cross zero and white otherwise. In all panels, light gray shading indicates quarters of regular-season flowering (first and second quarter), and blue indicates quarters with anomalous flowering (fourth quarter 2018 and 2025).

## Discussion

Some of the first observed, and best documented, impacts of climate change have been on the timing and intensity of seasonal ecological processes (Cleland et al., 2007; Piao et al., 2019). Despite substantial research documenting these impacts, we are aware of no study attempting to directly attribute specific extreme ecological events to global climate trends. Here, we make such an attempt for a recent dramatic, out-of-season flowering event in populations of Joshua tree. With extensive observation data provided by the iNaturalist crowdsourcing platform, we are able to train a machine learning model that identifies biologically plausible weather triggers of seasonal flowering, and which recovers flowering activity recorded in our training data and in independent records, as well as specific anomalous flowering events that motivated our study. However, comparing our model’s projections of flowering under actual conditions to projections under simulated counterfactual conditions without anthropogenic climate forcing, we find few significant differences in regular-season flowering activity, and none in quarters with anomalous flowering. This indicates that, despite prior work showing climate trends since the early 20th century have changed the frequency of Joshua tree masting years (Yoder et al., 2024), global climate change has not altered the frequency or intensity of out-of-season flowering in these species (Fig. 2D,E). By the same token, our results suggest winter flowering may be a natural response of Joshua tree populations to rare out-of-season precipitation events.

Attribution may be particularly difficult in this context, because climate models are notoriously poor at simulating historical sea surface temperature trends in the Pacific Ocean (Seager et al., 2019) and, by extension, hydroclimatic changes in the Western U.S. (Jacobson and Seager, 2025). While multidecadal precipitation changes in the western U.S. have been thought to be driven by natural climate variability (Lehner et al., 2018), there is emerging evidence that climate models may systematically underestimate the influence of climate change on Pacific Ocean variability and thus rainfall in the American west (Klavans et al., 2025). Further advances in physical climate science may be necessary to fully assess the effect of anthropogenic forcing on ecological systems governed by rainfall in this region. This work could plausibly recover more complex impacts of climate change on present-day phenology, or could find that such impacts will only emerge later in the century. Our study nevertheless shows that rising winter temperatures do not explain the anomalous winter blooms in 2018 and 2025—a rare exception to an extensive scientific literature on, and general public understanding of, the effects of climate change on plant phenology.

Our result also has immediate relevance for conservation of Joshua trees, which have received steadily increasing attention as an “umbrella species” for the Mojave Desert’s endemic biodiversity, and because of gathering evidence that projected climate change threatens their population viability (Smith et al., 2023). In addition to our work modeling the impacts of recent climate change on Joshua tree masting (Yoder et al., 2024), high-resolution species distribution modeling indicates that both *Y. brevifolia* and *Y. jaegeriana* stand to lose large fractions of their current suitable habitat under future climate scenarios (Shryock et al., 2025). Both Joshua tree species have been repeatedly considered for protection under the U.S. Endangered Species Act (U.S. Fish and Wildlife Service, 2023), and *Y. brevifolia* is protected under California’s Western Joshua Tree Conservation Act (WJT, 2023). Establishing that out-of-season flowering events represent low-frequency but not aberrant responses to seasonal variation in precipitation places them in the context of other risks facing these species as a result of climate change (Smith et al., 2023).

Across fields, climate change impact attribution studies are an area of rapidly growing interest and methodological innovation (Carleton and Hsiang, 2016; Carlson et al., 2024, 2025; Dudney et al., 2025). This study is among the first to show that existing platforms for biodiversity monitoring can support attribution work, even if most available data were collected in the last one to two decades. Future work should continue to develop models that can identify predictable and transferrable relationships between weather and important ecological patterns and processes; and it should continue to test the role of climate change in a broad range of phenomena, including those that may be the result of natural variability. In doing so, we can not only improve the overall evidence base for understanding climate change impacts on ecosystems, but also develop rapid attribution methods that can work on timescales aligned with public interest and ecosystem management needs.

## Acknowledgments

Support was provided by CSU Northridge (to JBY) and a Yale School of Public Health Transformation Award (to CJC). We thank the iNaturalist contributor community for making this work possible, particularly Drew Kaiser, who provided the image we use as Fig. 1A; the USA National Phenology Network and contributors to its *Nature’s Notebook* program; and Christopher I. Smith for helpful comments. Finally, we thank Alan Jay Lerner and Frederick Loewe, for their reminder that the only place where ecologically important seasonal events hold to a convenient, invariable schedule is in Camelot.

## Data accessibility

Code for the acquisition and management of training data, and for running all analyses, is available for review on GitHub: github.com/JBYoderLab/jotr_bonus_blooms

## Author contributions

All authors contributed to conceiving and designing the project, and to writing the manuscript; JBY curated iNaturalist records and managed model development with CJC; CWC simulated counterfactual conditions; JBY projected flowering activity under factual and counterfactual conditions and performed downstream analyses.

## Conflict of interest

The authors declare no conflict of interest.

## Supporting information for: Limited role of climate change in out-of-season Joshua tree flowering

**Fig. S1.**
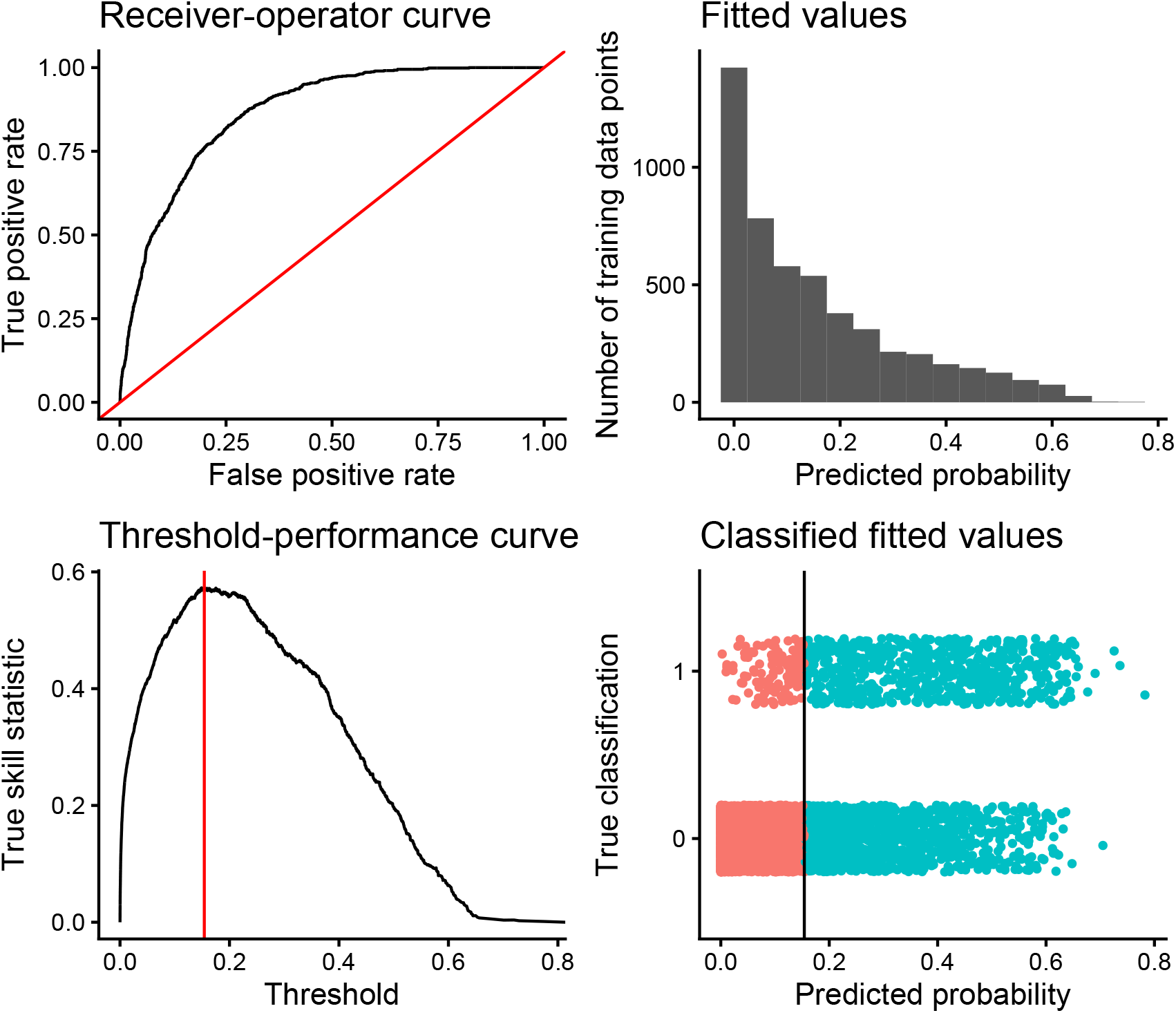
Performance and cutoff optimization for the RI-BART model, trained on 5 070 spatially dispersed records of quarterly flowering activity for both Joshua tree species, with the six predictors given in the main text and a RI effect representing quarter of observation. Top left: Receiver-operator curve for the model; *AUC* = 0.86. Top right: histogram of fitted values (probability of flowering) for the full training data set. Lower left: Threshold-performance curve, identifying a classification cutoff of 0.29 to maximize true positives while minimizing false positives. Lower right: Fitted values and classifications for the training data set.

**Fig. S2.**
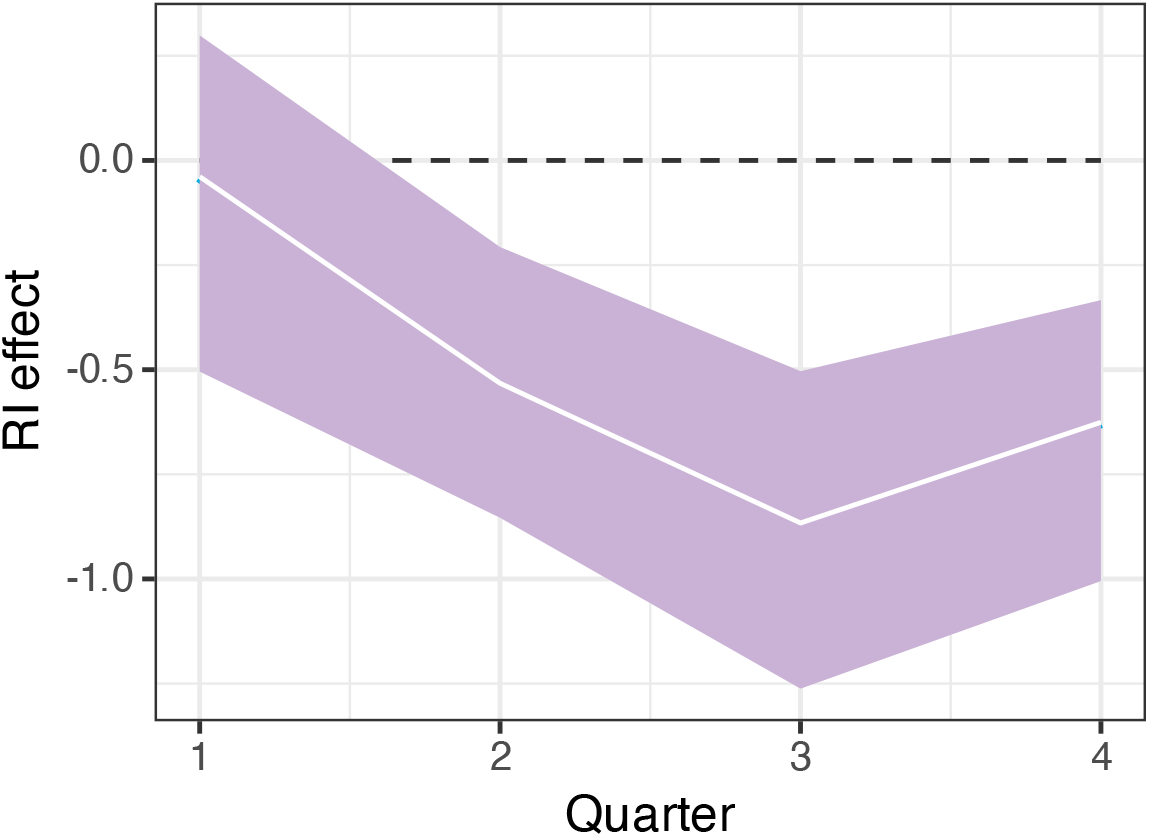
Plot of posterior effect estimates for the RI effect of quarter in our final working model.

**Fig. S3.**
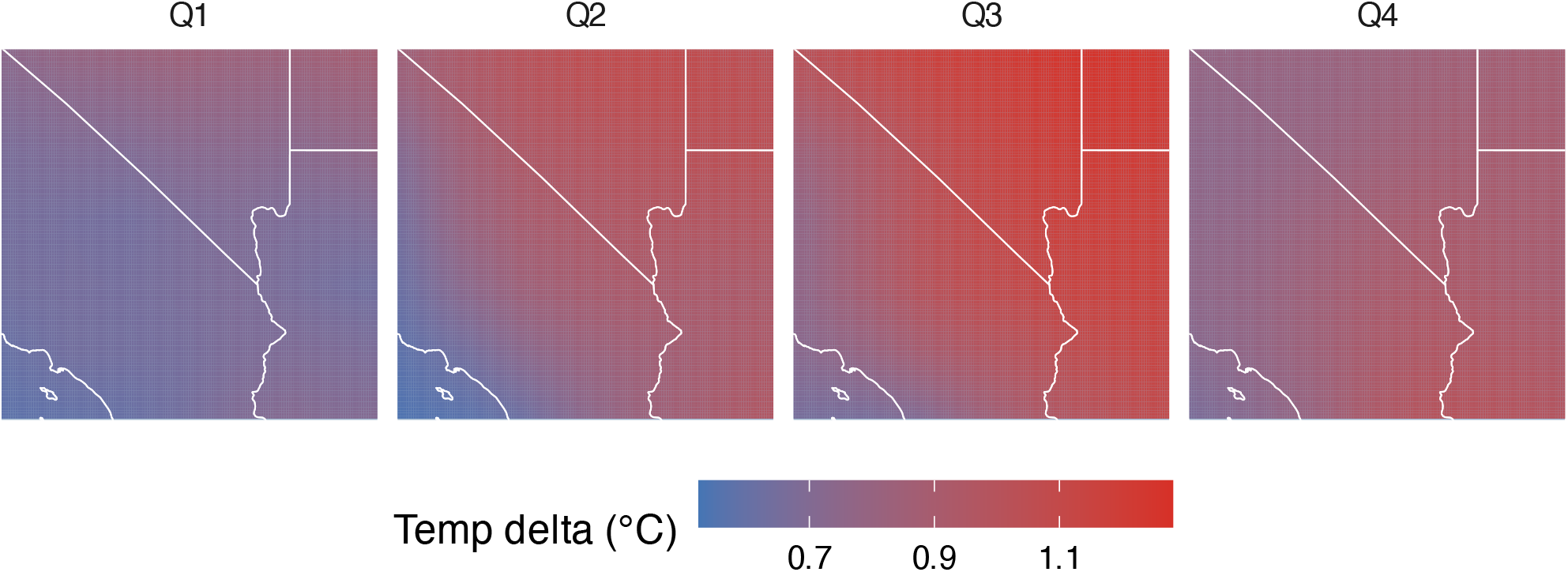
Temperature deltas for simulated counterfactual weather, by calendar quarter

**Fig. S4.**
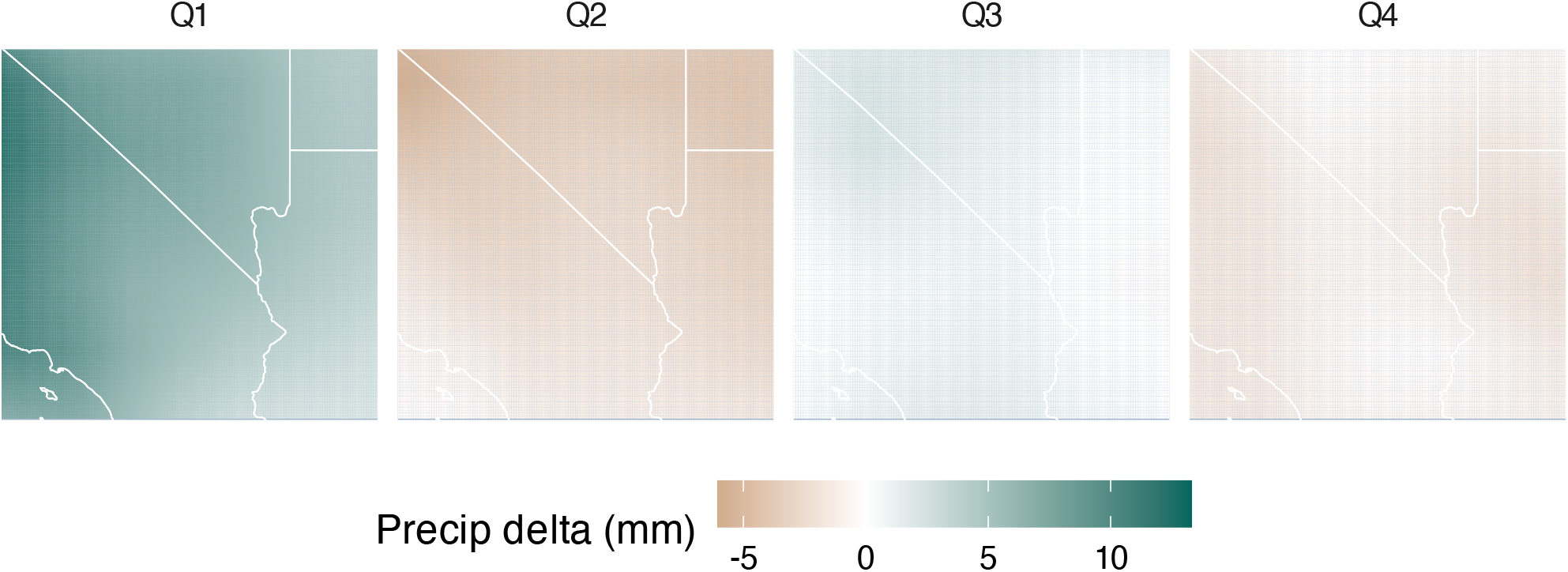
Precipitation deltas for simulated counterfactual weather, by calendar quarter

**Fig. S5.**
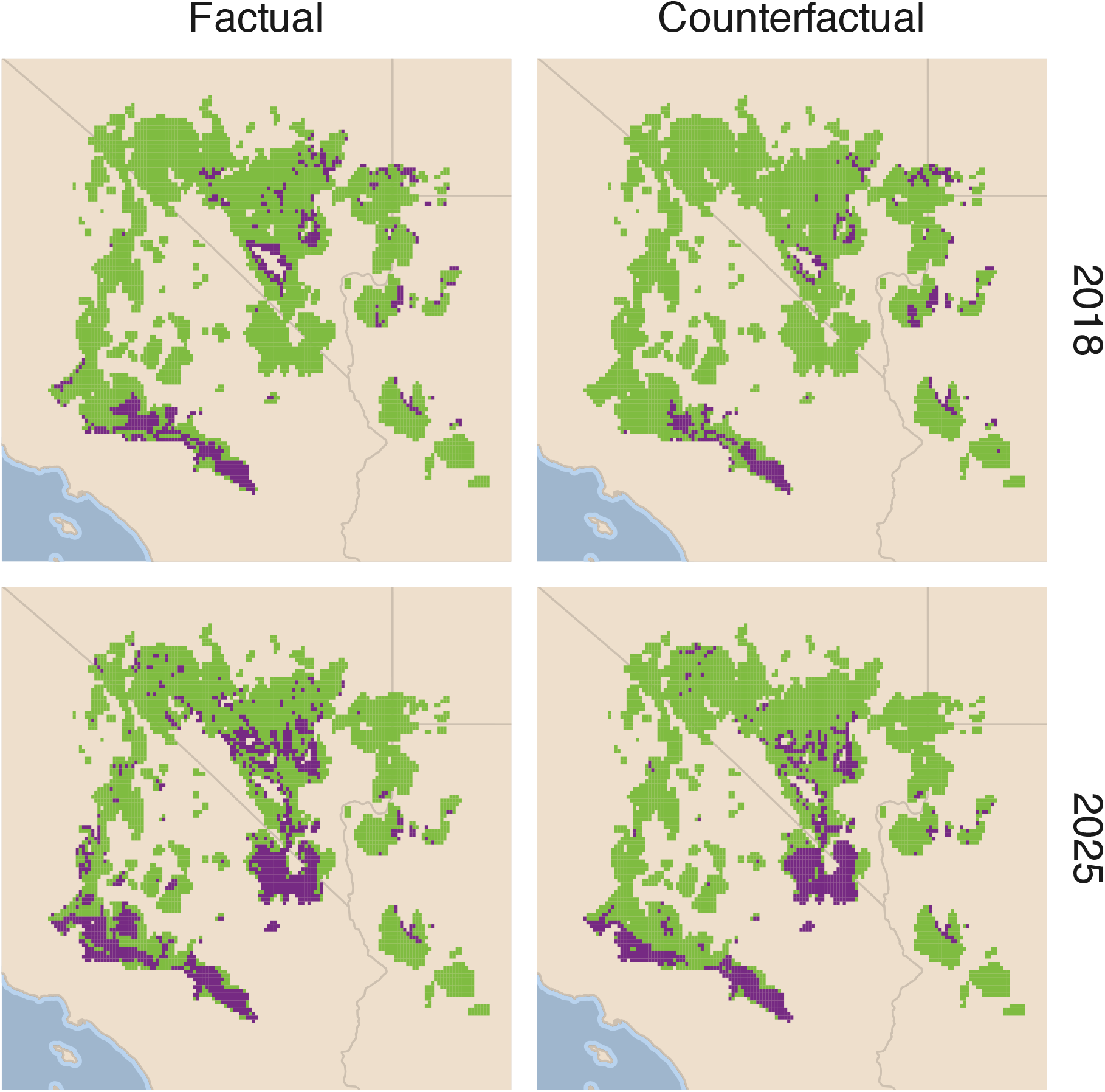
Extent of flowering (purple shading) in Q4 of 2018 and 2025, the two observed anomalous winter flowering events, as projected from factual weather data and simulated counterfactual conditions using our RI-BART model.

## Notes

https://github.com/JBYoderLab/jotr_bonus_blooms

### Competing Interest Statement

The authors have declared no competing interest.

### Summary of Updates

Revisions for journal submission; expanding Introduction and Discussion context

## Literature cited

2023. Western Joshua Tree Conservation Act, California Fish and Game Code Division 2, Chapter 11.5.

Bai, H., C. Strong, J. M. LaMontagne, I. V. Widick, and B. Zuckerberg. 2024. A North American climate-masting-irruption teleconnection and its change under future climate. Science of The Total Environment 948:174473.

Baum, J. K., M. A. Slein, J. C. Garen, Z. Sang, S. Emry, K. J. A. Goodwin, C. G. Collins, J. Lewthwaite, M. Tseng, S. T. Michaletz, A. C. Burton, C. D. G. Harley, D. S. Srivastava, The 2021 Western North American Heat Dome Data Consortium, P. K. Abram, A. L. Angert, E. Bayne, L. D. Daniels, S. Dickson-Hoyle, H. Earle, M. T. Franklin, A.-L. M. Gehman, T. Gharajehdaghipour, D. Hodder, L. Leston, A. McAfee, M. G. E. Mitchell, K. A. Otter, T. Richardson, J. Rosenfeld, G. Sadlier-Brown, P. Sandoval-Acuña, C. Shores, S. Starko, J. Thiessen, B. Timmer, K. Tjaden-McClement, S. Tseng, Y. Uriel, and B. William. 2026. Widespread ecological responses and cascading effects of the 2021 western North American heatwave. Nature Ecology & Evolution 10:864– 879.

Brenskelle, L., V. Barve, L. C. Majure, R. P. Guralnick, and D. Li. 2021. Analyzing a phenological anomaly in Yucca of the southwestern United States. Scientific Reports 11:20819.

California Consortium of Herbaria. 2024. CCH2 Portal. cch2.org/portal/index.php, accessed 12 Feb 2024.

Carleton, T. A., and S. M. Hsiang. 2016. Social and economic impacts of climate. Science 353:aad9837.

Carlson, C., D. Mitchell, T. Carleton, M. Chersich, R. Gibb, T. Lavelle, M. Lukas-Sithole, M. North, C. Lippi, M. New, S. Ryan, S. Shumba, and C. Trisos. 2024. Designing and describing climate change impact attribution studies: a guide to common approaches. EarthArXiv .

Carlson, C. J. 2020. embarcadero: Species distribution modelling with Bayesian additive regression trees in r. Methods in Ecology and Evolution 11:850–858.

Carlson, C. J., S. N. Bevins, and B. V. Schmid. 2022. Plague risk in the western United States over seven decades of environmental change. Global Change Biology 28:753–769.

Carlson, C. J., D. Mitchell, R. Gibb, R. F. Stuart-Smith, T. Carleton, T. E. Lavelle, C. A. Lippi, M. Lukas-Sithole, M. A. North, S. J. Ryan, et al. 2025. Health losses attributed to anthropogenic climate change. Nature Climate Change 15:1052–1055.

Chipman, H. A., E. I. George, and R. E. McCulloch. 2010. BART: Bayesian additive regression trees. The Annals of Applied Statistics 4:266 – 298.

Cleland, E. E., I. Chuine, A. Menzel, H. A. Mooney, and M. D. Schwartz. 2007. Shifting plant phenology in response to global change. Trends in Ecology & Evolution 22:357–365.

Diffenbaugh, N. S., F. V. Davenport, and M. Burke. 2021. Historical warming has increased US crop insurance losses. Environmental Research Letters 16:084025.

Dorie, V. 2023. dbarts: Discrete Bayesian additive regression trees sampler. R package version 0. 9-23. https://CRAN.R-project.org/package=dbarts.

Dudney, J., L. E. Dee, R. Heilmayr, J. Byrnes, and K. Siegel. 2025. A causal inference framework for climate change attribution in ecology. Ecology Letters 28:e70192.

Esque, T. C., D. F. Shryock, G. A. Berry, F. C. Chen, L. A. DeFalco, S. M. Lewicki, B. L. Cunningham, E. J. Gaylord, C. S. Poage, G. E. Gantz, R. A. Van Gaalen, B. O. Gottsacker, A. M. McDonald, J. B. Yoder, C. I. Smith, and K. E. Nussear. 2023. Unprecedented distribution data for Joshua trees (Yucca brevifolia and Y. jaegeriana) reveal contemporary climate associations of a Mojave Desert icon. Frontiers in Ecology and Evolution 11:1266892.

Eyring, V., S. Bony, G. A. Meehl, C. A. Senior, B. Stevens, R. J. Stouffer, and K. E. Taylor. 2016. Overview of the coupled model intercomparison project phase 6 (cmip6) experimental design and organization. Geoscientific Model Development 9:1937–1958.

Gillett, N. P., H. Shiogama, B. Funke, G. Hegerl, R. Knutti, K. Matthes, B. D. Santer, D. Stone, and C. Tebaldi. 2016. The detection and attribution model intercomparison project (DAMIP v1. 0) contribution to CMIP6. Geoscientific Model Development 9:3685–3697.

Godsoe, W., J. Yoder, C. Smith, and O. Pellmyr. 2008. Coevolution and divergence in the Joshua tree/yucca moth mutualism. The American Naturalist 171:816–823.

IPCC. 2021. Summary for policymakers. In V. Masson-Delmotte, P. Zhai, A. Pirani, S. L. Connors, C. Péan, S. Berger, N. Caud, Y. Chen, L. Goldfarb, M. I. Gomis, M. Huang, K. Leitzell, E. Lonnoy, J. B. R. Matthews, T. K. Maycock, T. Waterfield, O. Yelekçi, R. Yu, and B. Zhou, eds., Climate Change 2021: The Physical Science Basis. Contribution of Working Group I to the Sixth Assessment Report of the Intergovernmental Panel on Climate Change. Cambridge University Press, Cambridge, United Kingdom and New York, NY, USA.

Jacobson, T. W.-P., and R. Seager. 2025. Pacific decadal variability and its hydroclimate teleconnections in cmip6 models. Journal of Climate 38:5103–5127.

Klavans, J. M., P. N. DiNezio, A. C. Clement, C. Deser, T. M. Shanahan, and M. A. Cane. 2025. Human emissions drive recent trends in North Pacific climate variations. Nature 644:684–692.

Kuta, S. 2026. The American Southwest’s Iconic Joshua Trees Are Blooming Early—and Scientists Want Your Help to Figure Out Why. Section: Smart News, Smart News Science.

Lehner, F., C. Deser, I. R. Simpson, and L. Terray. 2018. Attributing the US Southwest’s recent shift into drier conditions. Geophysical Research Letters 45:6251–6261.

NASEM. 2016. Attribution of extreme weather events in the context of climate change. National Academies Press.

Parmesan, C., C. Duarte, E. Poloczanska, A. J. Richardson, and M. C. Singer. 2011. Overstretching attribution. Nature Climate Change 1:2–4.

Parmesan, C., M. Morecroft, Y. Trisurat, R. Adrian, G. Anshari, A. Arneth, Q. Gao, P. Gonzalez, R. Harris, J. Price, N. Stevens, and G. Talukdarr. 2022. Terrestrial and Fresh-water Ecosystems and Their Services. Cambridge University Press, Cmabridge, UK.

Parmesan, C., and G. Yohe. 2003. A globally coherent fingerprint of climate change impacts across natural systems. Nature 421:37–42.

Peattie, D. C. 1950. Joshuatree. Pages 303–309 in A Natural History of Western Trees. Bonanza Books, New York, NY.

Pellmyr, O. 2003. Yuccas, yucca moths, and coevolution: A review. Annals of the Missouri Botanical Garden 90:35.

Piao, S., Q. Liu, A. Chen, I. A. Janssens, Y. Fu, J. Dai, L. Liu, X. Lian, M. Shen, and X. Zhu. 2019. Plant phenology and global climate change: Current progresses and challenges. Global Change Biology 25:1922–1940.

PRISM Climate Group, Oregon State University. 2014. https://prism.oregonstate.edu.

R Core Team. 2025. R: A language and environment for statistical computing. R Foundation for Statistical Computing, Vienna, Austria. https://www.R-project.org.

Rode, E. 2026. Joshua trees are flowering in the Calif. desert. That’s bad news. SFGATE .

Seager, R., M. Cane, N. Henderson, D.-E. Lee, R. Abernathey, and H. Zhang. 2019. Strengthening tropical Pacific zonal sea surface temperature gradient consistent with rising greenhouse gases. Nature Climate Change 9:517–522.

Seneviratne, S., and X. Zhang. 2021. Weather and climate extreme events in a changing climate. Pages 1513–1766 in Climate Change 2021: The Physical Science Basis. Contribution of Working Group I to the Sixth Assessment Report of the Intergovernmental Panel on Climate Change. Cambridge University Press.

Shryock, D. F., T. C. Esque, G. A. Berry, and L. A. DeFalco. 2025. Assessing uncertainty in forecasts of refugia for Joshua trees using high-density distribution data. Ecosphere 16:e70308.

Smith, C. I., L. C. Sweet, J. Yoder, M. R. McKain, K. Heyduk, and C. Barrows. 2023. Dust storms ahead: Climate change, green energy development and endangered species in the Mojave Desert. Biological Conservation 277:109819.

Stager, M., P. M. Benham, N. R. Senner, R. R. Fitak, K. Denmead, R. K. Andringa, J. K. Grace, D. L. Dittmann, J. Siegrist, and A. M. Forsman. 2026. Storm-induced mass mortality results in both immediate and long-term consequences for a migratory songbird. Nature Ecology & Evolution 10:907–918.

Swain, D. L., D. Singh, D. Touma, and N. S. Diffenbaugh. 2020. Attributing extreme events to climate change: a new frontier in a warming world. One Earth 2:522–527.

Terrill, R. S., J. M. Maley, W. L. Tsai, K. B. Fistanic, R. J. Free-land, A. Franceschelli, B. Lewis-Smith, L. L. Lu, and J. B. Yoder. 2019. Tricolored blackbirds feeding in Joshua tree inflorescences. Western Birds 50:180–182.

Texas and Oklahoma Regional Consortium of Herbaria. 2024. TORCH Portal. portal.torcherbaria.org/portal, accessed 17 Feb 2024.

U.S. Fish and Wildlife Service. 2023. Species status assessment report for Joshua trees (Yucca brevifolia and Yucca jaegeriana).

USA National Phenology Network. 2026. Plant and animal phenology data. Data type: Status and intensity. 2011–2026 for region: 37.66476°, -112.1414°(UR); 33.63007°, -112.1474°(LL).

Yoder, J. B., A. K. Andrade, L. A. DeFalco, T. C. Esque, C. J. Carlson, D. F. Shryock, R. Yeager, and C. I. Smith. 2024. Reconstructing 120 years of climate change impacts on Joshua tree flowering. Ecology Letters 27:e14478.

Yoder, J. B., and C. I. Smith. 2024. Joshua trees, yucca moths, and a natural history of interdependence. In S. Khalsa and J. Harrower, eds., Desert Forest: Life with Joshua Trees. Inlandia Books, Riverside, CA.

Zwiers, F., and G. Hegerl. 2008. Attributing cause and effect. Nature 453:296–297.

